# Tumor landscape of epithelial ovarian cancer highlights that EGR1 drives tumor invasion at single-cell resolution

**DOI:** 10.1101/2022.07.26.501637

**Authors:** Yuanfu Zhang, Shu Sun, Yue Qi, Yifan Dai, Yangyang Hao, Mengyu Xin, Rongji Xu, Hongyan Chen, Xiaoting Wu, Qian Liu, Congcong Kong, Guangmei Zhang, Peng Wang, Qiuyan Guo

## Abstract

Identifying underlying molecular mechanisms and biomarkers of epithelial ovarian carcinoma (EOC) proliferation and metastasis remains challenging. Patients of EOC are usually diagnosed at an advanced stage and the availability of invasion-related targets is limited. Herein, we explored the single-cell RNA sequencing (scRNA-seq) dataset of EOC and defined tumor physiological reprograming compared to bulk RNA-seq. The energy metabolism and anti-apoptotic pathway was found as critical contributors to intratumor heterogeneity. Moreover, hypoxia, oxidative phosphorylation (OXPHOS) and glycolysis were positively correlated, which have biologically activity trajectories during epithelial mesenchymal transition (EMT). The HMGH1, EGR1 and RUNX1 were found to be critical inducers of the EMT process in EOC. Experimental validation revealed that suppressed EGR1 decreased the expression of FAS and HSPG2 and associating with EMT progression in EOC. In tumor microenvironment (TME), CAFs were found have significant contribution to tumor immune infiltration and metastasis and accumulation of CAFs was associated with poorer patient survival. In conclusion, physiological features and molecular mechanisms in the TME of EOC were revealed and provided effective targets for the suppression of tumor metastasis.

## Introduction

Epithelial ovarian cancer (EOC) has a superior lethality rate compared to other gynecologic cancers which have higher incidence (1). Advanced diagnosis when the EOC has spread to the abdominal cavity and upper abdominal organs is one of the main factors contributing to high mortality (2). Nearly 10 years, EOC treatments have been updated and more targeted therapies are on the horizon. For example, poly ADP-ribose polymerase (PARP) inhibitors (3, 4), immunotherapy and heated intraperitoneal chemotherapy (HIPEC) provide a prospect of improving the survival rate of EOC (5, 6). Therefore, early diagnosis strategies for EOC and treatment of tumor metastasis are important for mortality improvement. However, the understanding of the mechanisms of tumor cell metastasis in EOC patients is limited. Malignant cells together with other stromal cells such as cancer-associated fibroblasts (CAFs), mesenchymal, and immune cells etc., constitute the tumor microenvironment (TME). Together, the complex environment of these cells determines tumor progression. For example, mesenchymal cells and CAFs release signaling factors to assist in the metastasis and proliferation of malignant epithelial cells (7, 8). Endothelial cells provide blood vessels that can provide nutritional support for cancer cells in a hostile environment (9). All these suggest that each cell subset needs to perform its role based on specific physiological mechanisms (10), which is influenced by mutational and environmental factors including somatic driver mutation and tissue origin (11).

Further, each cell has specific physiological state for their specific environment including spatial location, nutrient supply and communication with surrounding cells (12-14). Thus, conventional bulk sequencing reflecting the average expression level of bulky tumors is not sufficient to reflect the specific performance of each cell and cell subset in the TME. Application of single-cell RNA sequencing (scRNA-seq) technology in EOC reveals tumor microenvironment map and cellular characteristics (15-17). Although these studies represent the cutting-edge progress of EOC in scRNA-seq, the cellular map and molecular mechanism landscape of EOC still need continuous improvement.

To determine the cellular compositional characteristics and developmental mechanisms of EOCs, we analyzed the scRNA-seq datasets including 2794 single cells from 7 patients with different tissue sites (18) and identified the mechanism of tumor cell metastasis and invasion mediated by EGR1.

## Results

### Single-cell landscapes demonstrate tumor heterogeneity due to CNVs

We performed a systemic pipeline to analyze the single-cell RNA-seq profiles of EOC (Figure 1A). The 2794 single cells were classified into 11 clusters using UMAP (20) (Figure 1B) and were annotated as seven cell types based on the cell markers (16) (Figure 1C; see Supplementary Method). Contact the patient for clinical information, we found that benign, low-grade and high-grade serous EOC cells in cluster 1 were clearly distributed in three regions. Moreover, multiple cell clusters originated from several patients were found in malignant epithelial, fibroblasts and mesenchymal cells. All these suggest that the tumor heterogeneity exists in EOC, which was probably caused by copy number variations (CNVs) (33); this hypothesis was confirmed by InferCNV. Clusters 0, 6 and 9 defined as epithelial cells have a global accumulation of CNVs (Supplementary Figure S2A), suggesting the malignancy of these subsets. Despite tumor heterogeneity, high-grade serous ovarian cancers possessed deletions in chromosomes 13 and amplifications in chromosomes 8. Furthermore, immune cells were the most abundant component in addition to malignant epithelial cells (Figure 1D), where the immune cells content of benign, low-grade and high-grade serous ovarian cancers increased sequentially. The overall proportion of immune cells in each clinical subtype of tumor may reflect the different efficiencies of immune depletion (16), which may be a plausible explanation for why high-grade serous ovarian cancers is more suitable for immune-targeted therapy (34). Immune cells and malignant epithelial cells originating from different patients were distributed in separate clusters, while this phenomenon was not observed in stromal cells (Supplementary Figure S2B and C). These results further confirmed that tumor heterogeneity and stromal cell unity.

**Figure 1.**
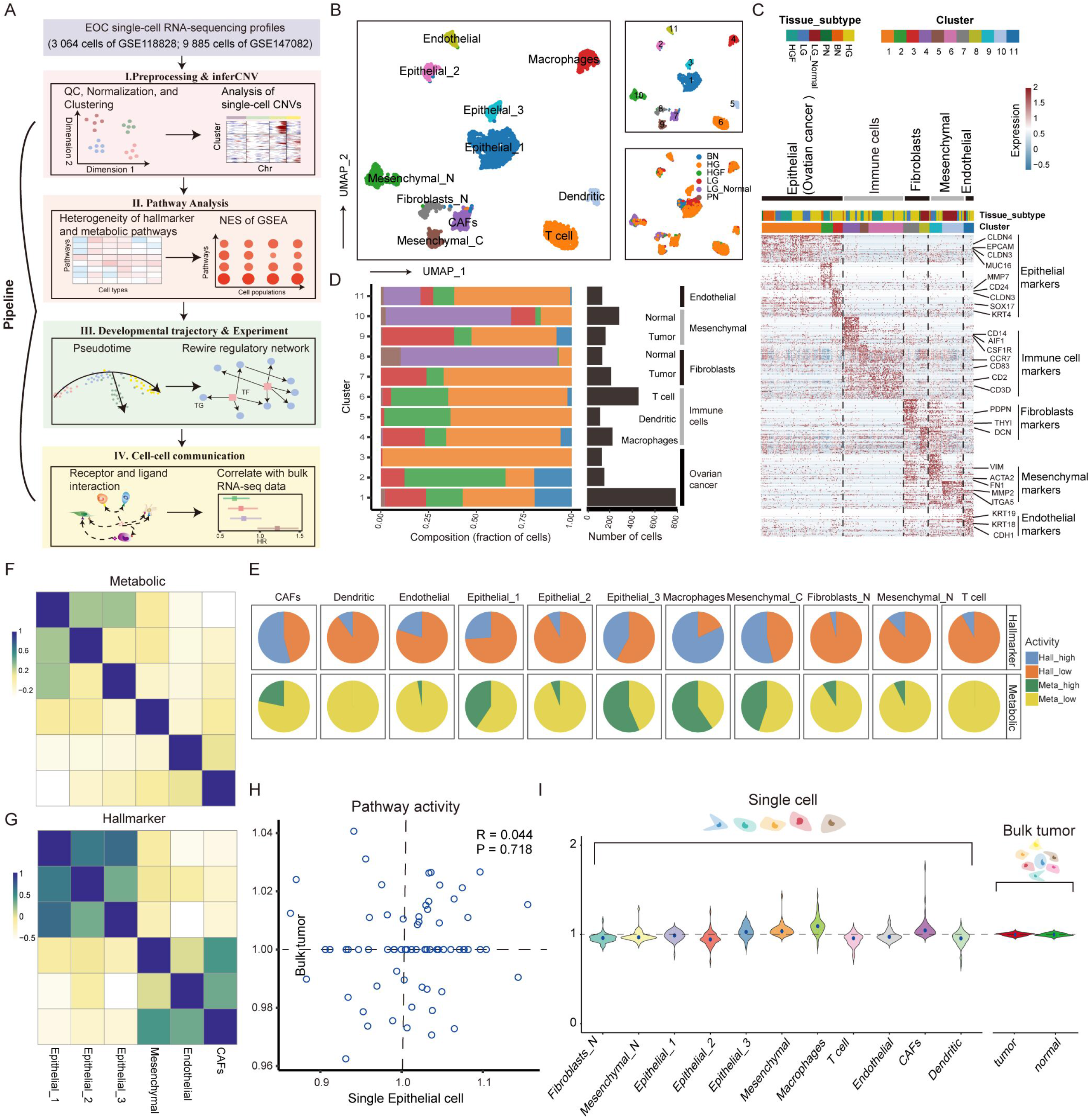
Single cell landscape of EOC. (A) Diagram of the multiple components and workflows of pipeline. (B) UMAP dimensionality reduction plot show cell type annotations, clustering results including 11 clusters, and clinical typing of cells. (C) Heatmap of differentially expressed genes. Expression levels of the top 50 genes (rows) differentially expressed in each cluster (column). (D) The number of cells per cluster and total number of cells in each clinical subtype, which are distributed among the identified cell types. (E) Proportion of up- and down-regulated pathways in metabolic pathways and hallmark among cell types. (F) Correlation of cell types other than immune cells in metabolic pathway activity. (G) Correlation of cell types other than immune cells in hallmark gene sets. (H) Scatter plot comparing pathway activity between OV bulk tumors in GEO and individual malignant cells in the scRNA-seq dataset. (I) The left panel shows the distribution of pathway activity in different cell types in single-cell RNA-seq, and the right panel shows the pathway activity in bulk tumor and normal samples from GEO.

### Cell type-specific reprograming of metabolic pathway and hallmarks

Tumorigenesis is a progress of malignant transformation of cell physiological functions including metabolism, signal transduction and cell proliferation and apoptosis (35). Therefore, mining the physiological characteristics of TME components can reflect the reprogramming of tumor ecology. The pathway activity evaluation algorithm (36) (see Supplementary Method) was used to calculate the pathway score of each cell type. There were 69 metabolic pathways which were found to be significantly associated with different EOC cell clusters. More than 60 pathways were highly activated (activity score >1 and p-value < 0.01) in at least one cell type (Supplementary Figure S3A). Similarly, there were 50 cancer hallmark genes sets which were highly activated in at least one cell type (Supplementary Figure S3B). As an example, the different activity of oxidative phosphorylation (OXPHOS), glycolysis, TCA cycle, and hypoxia in cell types was supported by direct comparison of the average expression of these pathways in each cell type (one-way ANOVA *p* < 0.01, Supplementary Figure S4). Compared with other cells, macrophages cells were significantly associating with more positive activation of metabolic pathways and cancer hallmarks (Figure 1E). These up-regulated gene sets were included in several parts of physiological functions, such as OXPHOS, glycolysis and EMT. Macrophages cells have a global up-regulation of metabolic pathway activity compared to malignant epithelial cells, which is not observed in other cancer (36). Meanwhile, the heterogeneity of malignant epithelial cells reflected in metabolic activity and tumor proliferation and invasion mechanisms, where type-1 epithelial cell was hyper-metabolically active and higher EMT active, type-2 was meso-metabolically active, and type-3 was hypo-metabolically active (Supplementary Figure S3 and S5). Although the metabolic pathway activity was strongly influenced by cell subsets (Figure 1F), there was a high correlation in oncogenic function between the three malignant epithelial cells (Figure 1G). This pattern also was found in CAFs and mesenchymal cells. These results suggest that heterogeneity of tumor metabolic reprogramming does not largely influence oncogenic function.

Contrasting TME physiological features observed at single-cell and bulk resolution will contribute to the correction of EOC prior knowledge. The pathway activity scores were calculated based on microarray data for bulk tumor samples obtained from GEO (GSE26712) and was compared with the results detected by scRNA-seq profiles. We found that only 26 metabolic pathways and 14 hallmarks were up-regulated in tumor samples compared to normal samples (p-value < 0.01, Supplementary Figure S6 A and B). The correlation of physiological pathway activity between bulk tumor samples and malignant epithelial cells was not significant (Pearson’s R = 0.044, p-value = 0.718, Figure 1H, Supplementary Figure S6 C and D), suggesting expression specificity between cell types was masked by the fact that bulk data measure the average expression levels over a mixture of multiple cell types. Moreover, the variation in the distribution of pathway activity for single-cell was greater than bulk data (average standard deviation of pathway activity = 0.075 for single cells compared to 0.015 for bulk tumor, Figure 1I). Taken together, the programming of cellular-specific physiological functional activity can be better reflected at single-cell resolution than bulk resolution.

### Intratumor heterogeneity of EOC depend on energy metabolism

For malignant epithelial cells, which have tumor heterogeneity reflected in molecular mechanisms, we performed re-cluster analysis and eight clusters were identified (Figure 2A) that predominantly from seven patients, indicating that individual differences in tumor cells is evident. The phenomenon may be caused by the patient’s unique physiological environment including location-specific and nutrient supply (36). There is also an undefined cluster of cells overlapping with type 2 epithelial cells, suggesting that hypo-metabolically active malignant cells are widely distributed and physiologically similar between patients.

**Figure 2.**
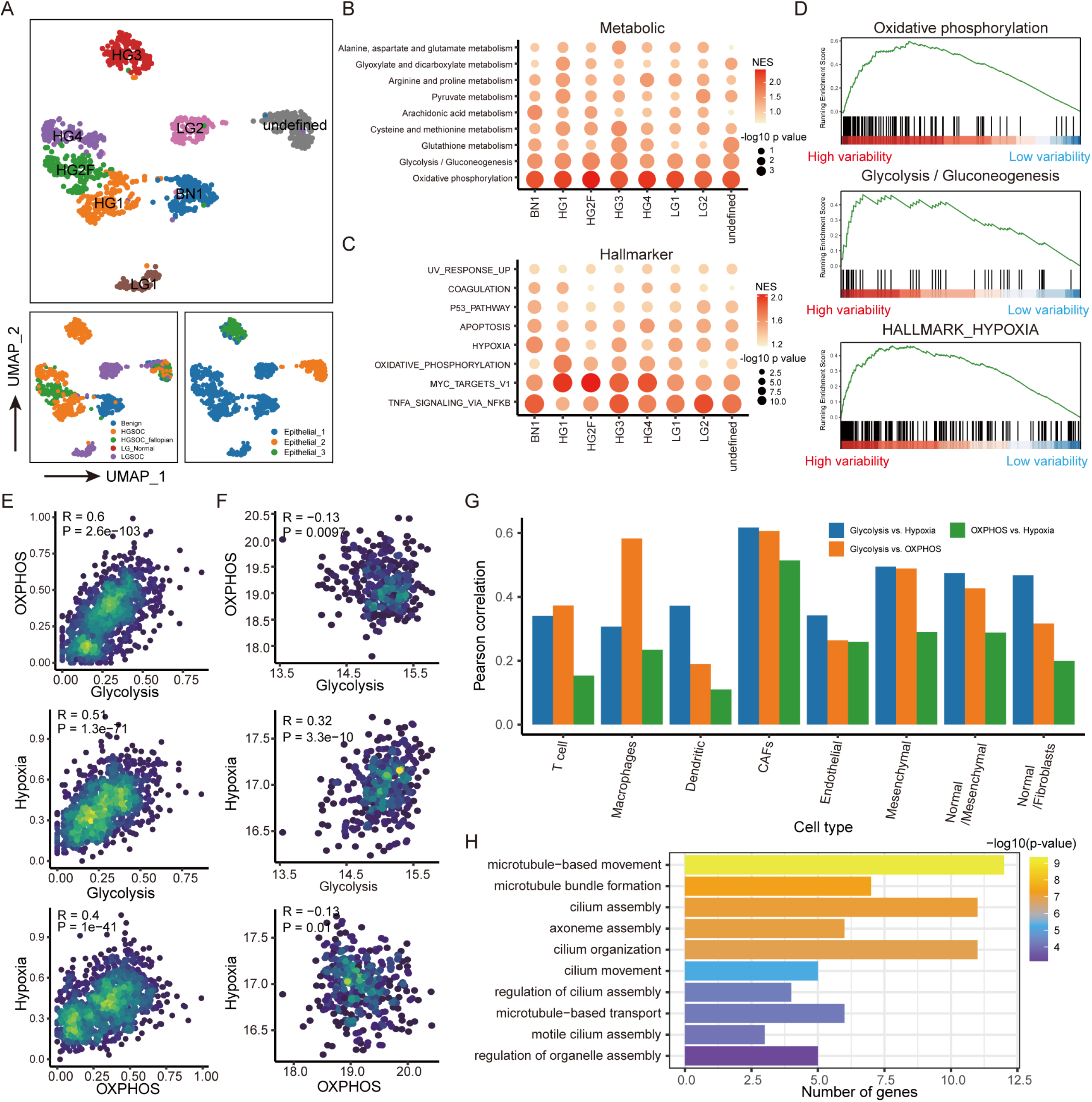
Major contributors to intratumor heterogeneity. (A) UMAP plots of malignant epithelial cell re-clustering. Each cell is labeled with tissue origin, clinical typing and first clustering result respectively. (B) The scatter plot demonstrating GSEA results for metabolic pathways weighted by principal component analysis. (C) The scatter plot demonstrating GSEA enrichment results for hallmark gene sets. (D) Enrichment of OXPHOS, glycolytic and hypoxic pathways on differential genes, which could explain most of the variation in the expression data. (E-F) Density scatter plot demonstrate glycolysis, OXPHOS activity and response to hypoxia in malignant epithelial cells based on single-cell and bulk data, respectively. The color of the dots indicates the local density. (G) Pearson correlation between glycolysis, OXPHOS activity and response to hypoxia in other cell type. (H) Enrichment of up-regulated genes in GO from low activity cells in OXPHOS, glycolytic and hypoxic pathway.

Reprogramming of cellular physiological mechanisms is largely determined by variation and space environment. To explore what aspects of cellular physiological mechanisms are affected by environmental factors. GSEA (23) was used to identify metabolic and cancer hallmarks enriched in genes that represents intratumor heterogeneity influenced by environment, mutation, etc. (Supplementary Figure S7). We found that the OXPHOS and NFKB signaling pathway had the highest enrichment scores in most tumor clusters (Figure 2B and C). Similarly, glycolytic and hypoxic signaling pathways also have potential contribution to intratumor heterogeneity across these tumor clusters. These results suggest that energy metabolism and cellular resistance to apoptosis were major contributors to intratumor heterogeneity of malignant epithelial cells. The coefficient of variation (CV), standard deviation (SD), and information entropy was then used as weights for each gene accounting for variation between malignant cells to exclude potential prejudice for formula selection and found consistent result in identification of functional pathways (Supplementary Figure S8).

Furthermore, the association between energy metabolism and environmental factors such as oxygen supply was performed. The hypoxic signals (hypoxia inducible factor, HIF) that indirectly reflect the oxygen content of cells were evaluated (Figure 2D). We found the high correlation between glycolysis and hypoxia (Pearson’ R = 0.51 for epithelial cells, Figure 2E), which is consistent with previous studies demonstrating that hypoxia activates glycolysis (37, 38). Moreover, the activity of OXPHOS was significantly correlated with glycolysis (Pearson’ R = 0.62) and the response to hypoxia (Pearson’ R = 0.43). Notably, we did not detect strong correlation between OXPHOS and glycolysis (Pearson’ R = -0.13) and between OXPHOS and hypoxia (Pearson’ R = -0.13) using bulk RNA-seq data of TCGA (Figure 2F). In addition to malignant epithelial cells, we had also identified the relationship between OXPHOS, glycolysis and hypoxia in stromal cells (Figure 2G), indicating the particular relationship between energy metabolism and hypoxia at single-cell resolution. For cells with low activity in energy metabolism, the up-regulated genes were significantly enriched in microtubule-based movement (Figure 2H), suggesting the subpopulation of malignant epithelial cells promote tumor progression (39). Taken together, energy metabolism major contributed to intratumor heterogeneity.

### Reprogramming of energy metabolism during EMT

We revealed that the higher metabolism activity of malignant cell had higher EMT score above. However, the relationship between energy metabolism, an important contributor to tumor heterogeneity, and tumor metastasis remains unknown. Further, the pseudo-developmental trajectory of epithelial cells, mesenchymal were simulated using monocle (Figure 3A). The EMT-cells were grouped into three branches (defined as “B1”, “B2” and “B3”) with different states. We found that the three epithelial cell subtypes were mainly concentrated in state1 and state3, while mesenchymal cells and CAFs were concentrated in state2 (Figure 3B). The EMT pathway activity gradually increased with pseudo-time development and reached the highest at state 2 (Figure 3C). The phenomenon was also observed in independent dataset (GSE147082) (Supplementary Figure S9A-E), indicating the pathology of epithelial cells transformation to mesenchymal cells. We found that the score of OXPHOS, glycolysis and hypoxia increased from branch 1 to branch 2 during EMT (Figure 3D-F), which is consistent with previous studies suggesting that reduced energy metabolism provide environment for EMT and hypoxic conditions promote the EMT (40, 41). Several branching-dependent genes were identified from branch1 to branch2 and most of them related to EMT pathway and energy metabolism (Figure 3G and Supplementary Figure S9I). Moreover, cell invasiveness and stemness follow similar trends to energy metabolism during EMT (Figure 3H and I). The OXPHOS and glycolysis were higher correlated with cell invasiveness (Pearson’s = 0.59 for OXPHOS, Pearson’s = 0.56 for glycolysis) and cell stemness (Pearson’s = 0.37 for OXPHOS, Pearson’s = 0.26 for glycolysis, Figure 3J). The high correlation between hypoxia and EMT (Pearson’s = 0.65) confirms the reliability of this study. Besides, 810 EMT-related genes (586 positively and 224 negatively; |R|>0.2 (25)) was identified based on pseudo time. GO functional enrichment results showed that positive EMT-related genes act mainly in the extracellular matrix and cell adhesion that may provide the requirements for cell transfer (Figure 3K) and negative EMT-related genes are associated with the encoding of membrane proteins (Figure 3L). These results indicate that hypoxia induces shift in the function of epithelial cells to promote tumor cell migration.

**Figure 3.**
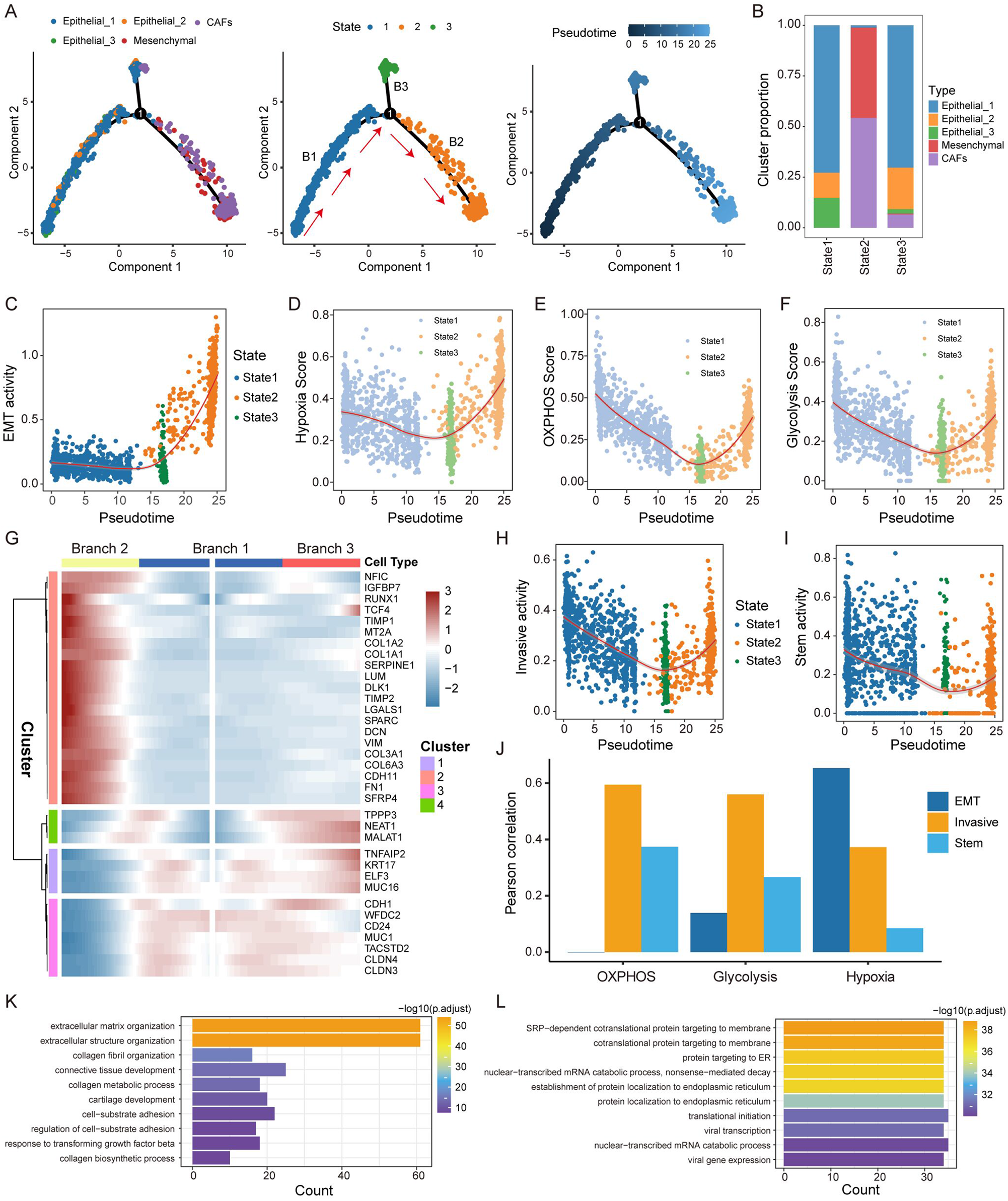
Specific energy supply during EMT. (A) Developmental trajectory of cells in two dimensions. Cells are colored according to cell type (left), state (middle), and pseudo-time (right). The red arrows indicate the defined “epithelial-mesenchymal path”. (B) The number of cells per cell types in each developmental state. (C) Variations in pathway activity of EMT with pseudo-time. (D-F) The pathway activity of OXPHOS, glycolytic and hypoxic distribution with pseudo-time. (G) The heatmap shows the branch-dependent genes at branch point 1. The center of the heatmap indicates branch B1, the left to B2 and the right indicates B3. (H) The invasive activity distribution with pseudo-time. (I) The stem activity distribution with pseudo-time. (J) The bar plot compared the pathway activity of OXPHOS, glycolytic and hypoxic with that of EMT, invasive and stem, respectively. (K-L) Enrichment of up-and down-regulated genes associated with the pseudo-time in the GO term.

### Prognostic marker EGR1 drives EMT in EOC

The driver genes of EMT related to pseudo-time were identified based on molecular regulatory network. In transcriptional regulatory network, we identified 41 positive-TFs and 10 negative-TFs from EMT-related genes (Figure 4A), which targeted 50 positive-target genes and two negative-target genes (Figure 4B, Supplementary Figure S9G). We found that FOSB, RUNX1, as important drivers of tumor proliferation and metastasis (42, 43), were significantly associated with OS of EOC patients (Figure 4C and D) and up-regulated during EMT (Figure 5F-H). It has been demonstrated that EGR1 can regulate angiogenesis to promote tumor cell metastasis (44), which up-regulated in the EMT process and regulated target-gene HSPG2 (Figure 5F-J, Supplementary Figure S9F-H) to enhanced EMT in this study. Intriguingly, high expression of both EGR1 and HSPG2 were associated with poorer OS in EOC patients (Figure 5E, Supplementary Figure S10A). Besides, the HMGA1 down-regulated expression of CDH1 encoding E-cadherin (Supplementary Figure S10B). Furthermore, the FOSB, RUNX1, HSPG2, and EGR1 were significantly different expressed between invasive epithelial (iE) and invasive mesenchymal (iM) samples defined from TCGA-OV data (45, 46) (Figure 5K and Supplementary Figure S11), suggesting that several critical genes driving the EMT process can only be identified at single-cell resolution.

**Figure 4.**
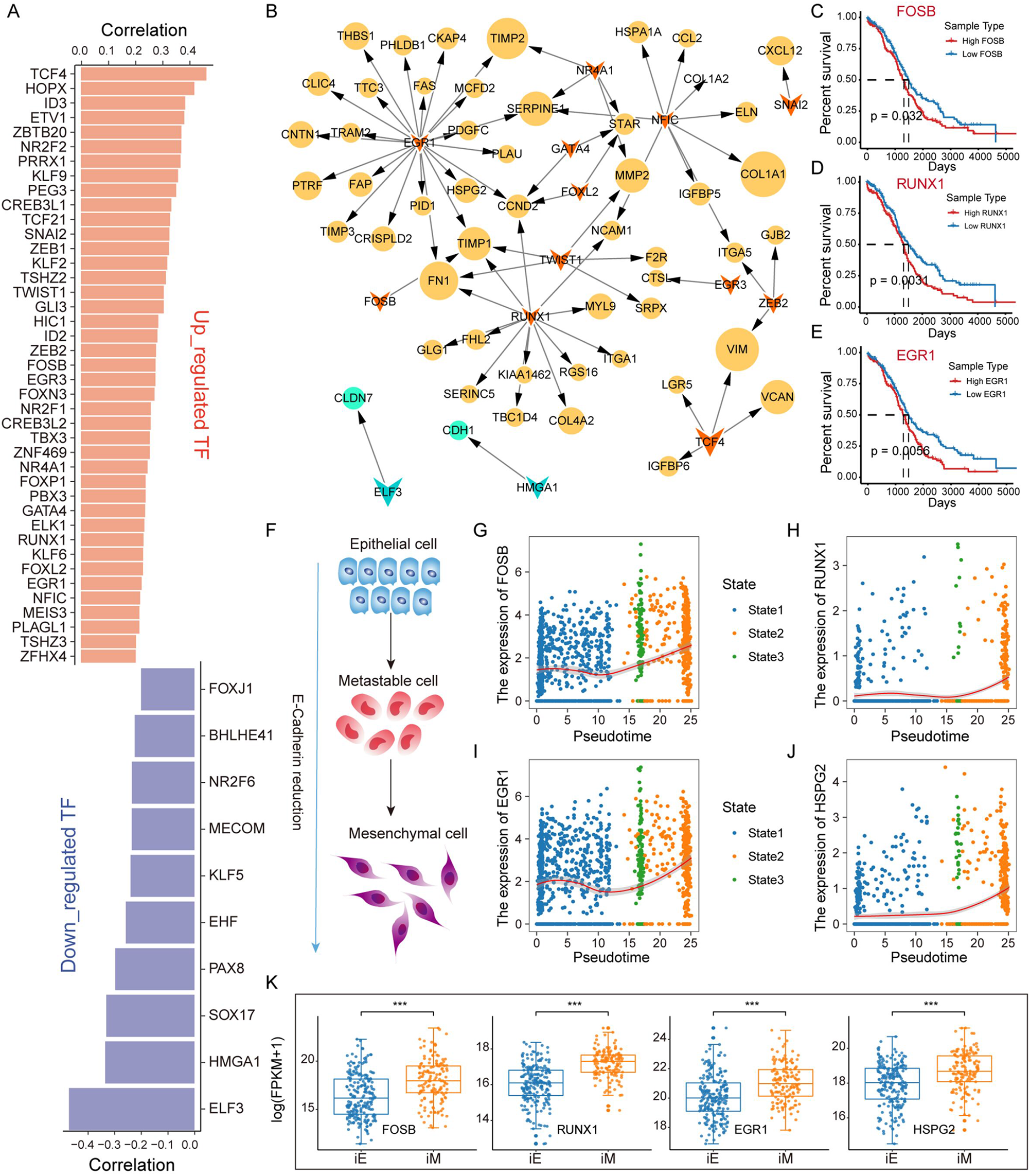
Critical factors boost EMT in EOC. (A) Up- and down-regulated TFs and their Pearson correlation coefficients with pseudo-time. (B) Transcriptional regulatory network constructed using temporal-associated genes. The diamond represents TF and the circle represents target gene. The size of the graph is determined by the coefficients of positively (yellow) and negatively (green) genes. (C-E) Survival curves of FOSB, RUNX1 and EGR1 gene expression in TCGA-OV dataset. (F) Schematic diagram of the reversal of epithelial cells with reduced E-cadherin into mesenchymal cells. (G-J) Variation of FOSB, RUNX1, EGR1, and HSPG2, which are transcription factors and target genes, expression levels with pseudo-time. (K) Box plot of genes including FOSB, RUNX1, EGR1, and HSPG2 expression variations between iE and iM samples of bulk RNA-seq from TCGA.

**Figure 5.**
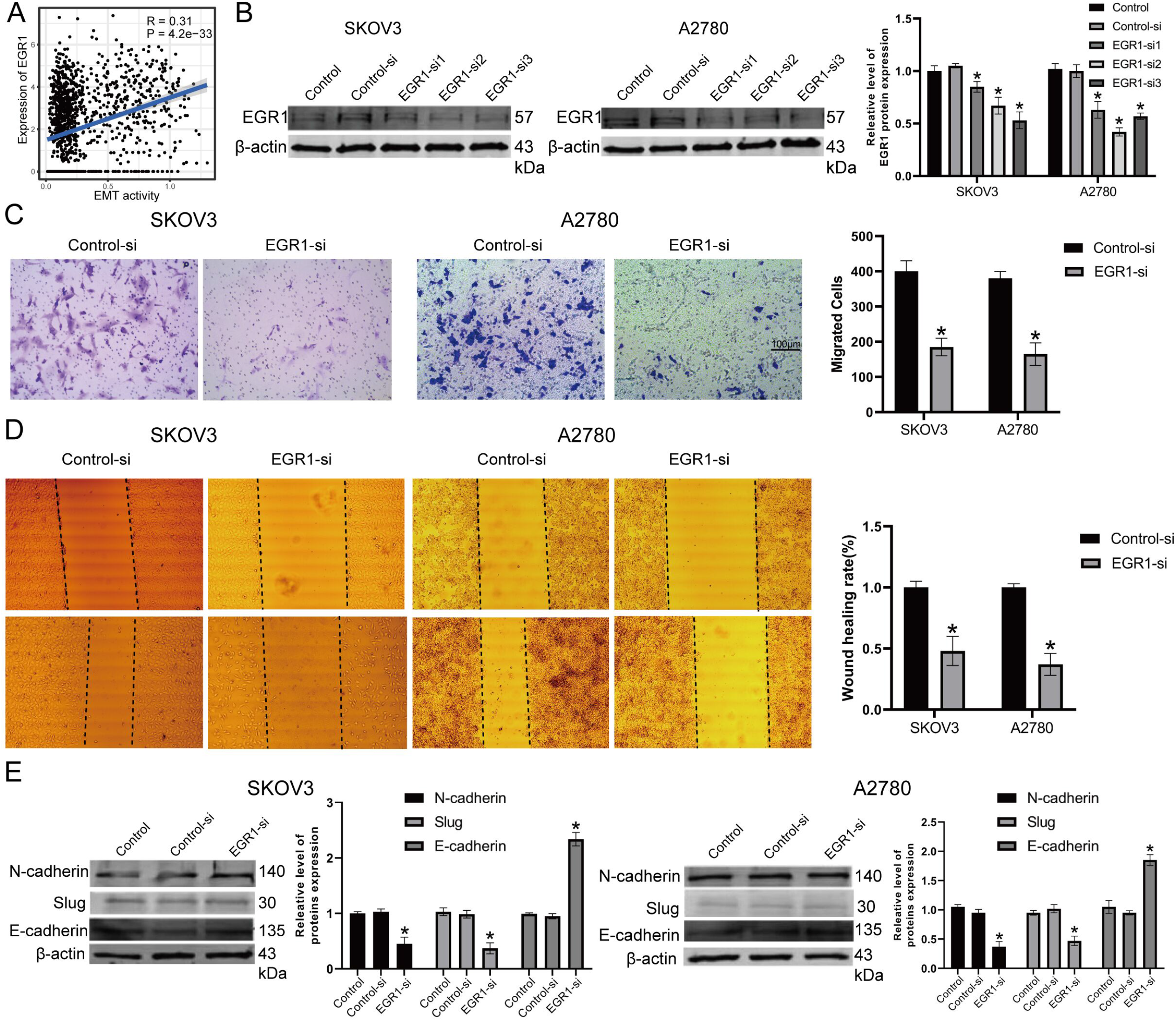
Knockdown EGR1 inhibit ovarian cancer cell migration and invasion. (A) Positive correlation was also observed between EMT activity and EGR1 expression in GSE118828 dataset. (B) Knockout efficiency of EGR1 in SKOV3 and A2780 cell lines. (C) Evaluation of EGR1 effect on the migration and invasion of ovarian cancer cells by transwell analysis. (D) Evaluation of EGR1 effect on the migration and invasion of ovarian cancer cells by wound healing analysis. (E) Significantly increased expression of E-cadherin and decreased expression N-cadherin and Slug were observed after EGR1 silencing.

### Knockdown EGR1 inhibit ovarian cancer cell migration and invasion

The TF EGR1 regulated the most EMT-related genes (Figure 4B) and exhibited significant prognostic efficiency of patient survival (Figure 4E). Positive correlation was also observed between EMT activity and EGR1 expression (Figure 4C, Figure 5A and Supplementary Figure S9J). Furthermore, experimental analysis by employing two ovarian cancer cell lines SKOV3 and A2780. Western blotting was used to show the Knockout efficiency of EGR1 in SKOV3 and A2780 (Figure 5B). We evaluated the effect of EGR1 on the migration and invasion of EOC cells by transwell and wound healing (Figure 5C-D). Migration and invasion capacity of SKOV3 and A2780 cells were significantly decreased compared to the control group, suggesting that the recovery of epithelial phenotype inhibits cell migration. On the other hand, we test the EMT-relative proteins in SKOV3 and A2780 after treatment with EGR1 inhibition (Figure 5E). Our data showed that silencing of EGR1 significantly increased the expression of E-cadherin. While the protein of N-cadherin and Slug were down-regulated in EGR1 group. These results indicate that silence of EGR1 suppressed migration and invasion in ovarian cancer cells.

### Suppressed EGR1 Decreases Expression of FAS and HSPG2

Next, the regulating mechanism of EGR1 was verified. The expression of two EGR1 target genes, FAS and HSPG2, were positively correlating with EGR1 based on the single cell expression profile and TCGA bulk dataset (Figure 6A-D). Within the TME of EOC, cells with high FAS and HSPG2 expression level exhibiting increased EMT activity (Figure 6E-F). To further confirm the transcriptional relationship between EGR1 and downstream target genes, we explored Chromatin Immunoprecipitation Sequencing (ChIP-seq) experiment dataset from ENCODE. Enriched EGR1 sequencing read peaks were found in the FAS and HSPG2 transcriptional factor binding region across different bio-samples, indicating directly transcriptional relationship between EGR1 and its targets (Figure 6G-H). The experiment demonstrated that FAS and HSPG2 expression at the protein level was markedly down-regulated in EGR1-si group in SKOV3 and A2780 cells compared with NC-si group (Figure 6I-J). These results indicated that EGR1 decreases the expression of FAS and HSPG2 and associating with EMT in the TME of EOC.

**Figure 6.**
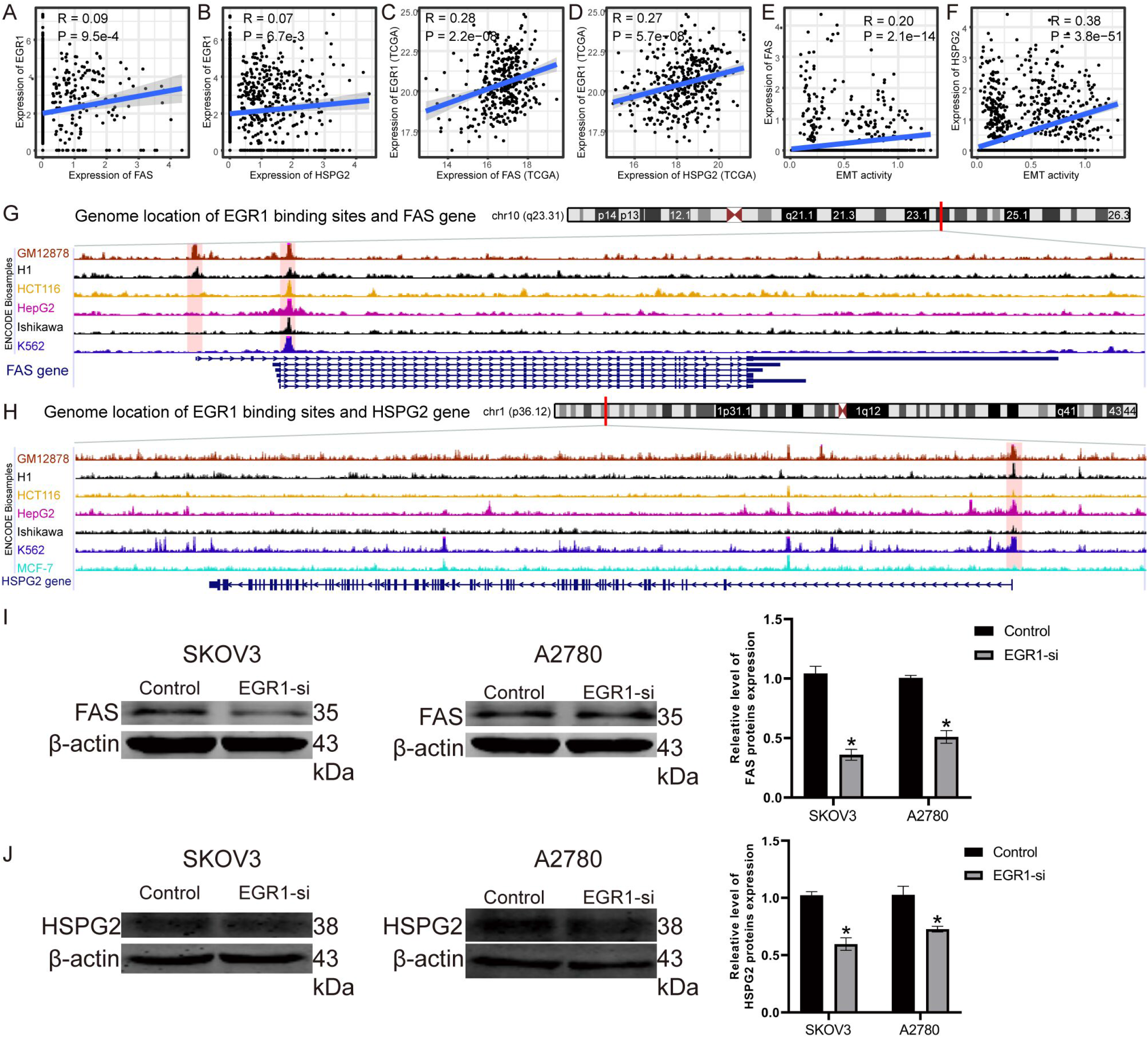
Suppressed EGR1 Decreases Expression of FAS and HSPG2. (A-D) Positive correlation was observed between EGR1 and its targets FAS and HSPG2 in GSE118828 and TCGA dataset. (E-F) The FAS and HSPG2 expression level were positively correlated with EMT activity. (G-H) Enriched EGR1 sequencing read peaks were found in the FAS and HSPG2 transcriptional factor binding regions. (I-J) The protein expression level of FAS and HSPG2 was markedly down-regulated in EGR1-si group in SKOV3 and A2780 cells.

### Cell-cell communication reveals the pro-metastatic effect of CAFs

The 42 signaling pathways was identified by CellChat (29) that enable cell-cell communication between nine cell subsets in tumor tissue, and the amount of signal emission in the three epithelial cell clusters correlated with metabolic pathway activity (Supplementary Figure S12A). Notably, CAFs cells exhibited most communication relationships with other cell types and have the strongest interactions with mesenchymal and macrophage cells (Figure 7A). To explore the functions of these signaling pathways, we clustered the above 42 signaling pathways into three categories related to growth factors and metastatic determined by their functions (Figure 7B). Further, we collected CCL and CXCL family ligand-receptor interaction pairs associated with tumor metastasis (Figure 7C-E). CAFs highly expressed the ligand CXCL12 and their actionable receptors including CXCR4 and ACKR3 (Figure 7C). We found that CXCR4 was highly expressed in immune cells such as dendritic, T cells, and macrophages and thus CAFs induced immune cells to control immune infiltration (47). CXCL12 on CAFs could also combined with CXCR4 in epithelial cells to induce EMT (8). CCL5 and CXCL2, secreted by CAFs, could interacted with ACKR1 that is highly expressed on the surface of endothelial cells (Figure 7C and F). This signaling pathway is associated with tumor metastasis and invasion (48, 49), which also identified in EOC and we pinpointed their derived from CAFs. The role of CAFs in tumor proliferation and metastasis revealed that EOC patients have individual clinical phenotypes due to the cellular composition of the tumor tissue. The fraction of the nine cell subsets in the OV samples from TCGA was calculated using cibersortX (Figure 7G). Correlating the cell fraction data with clinical information, we noted that part of stage III patients had higher fraction of CAFs but lower fraction of immune cells. The accumulation of CAFs was also associated with poorer OS, suggesting that CAFs plays an important role in tumor progression (Figure 7H and I). However, the accumulation of mesenchymal and dendritic cells was however associated with better OS, suggesting that they may be protective factors for EOC (Figure 7J, Supplementary Figure S12B). Taken together, CAFs were highlighted to promote EOC progression through the CXCL signaling pathway.

**Figure 7.**
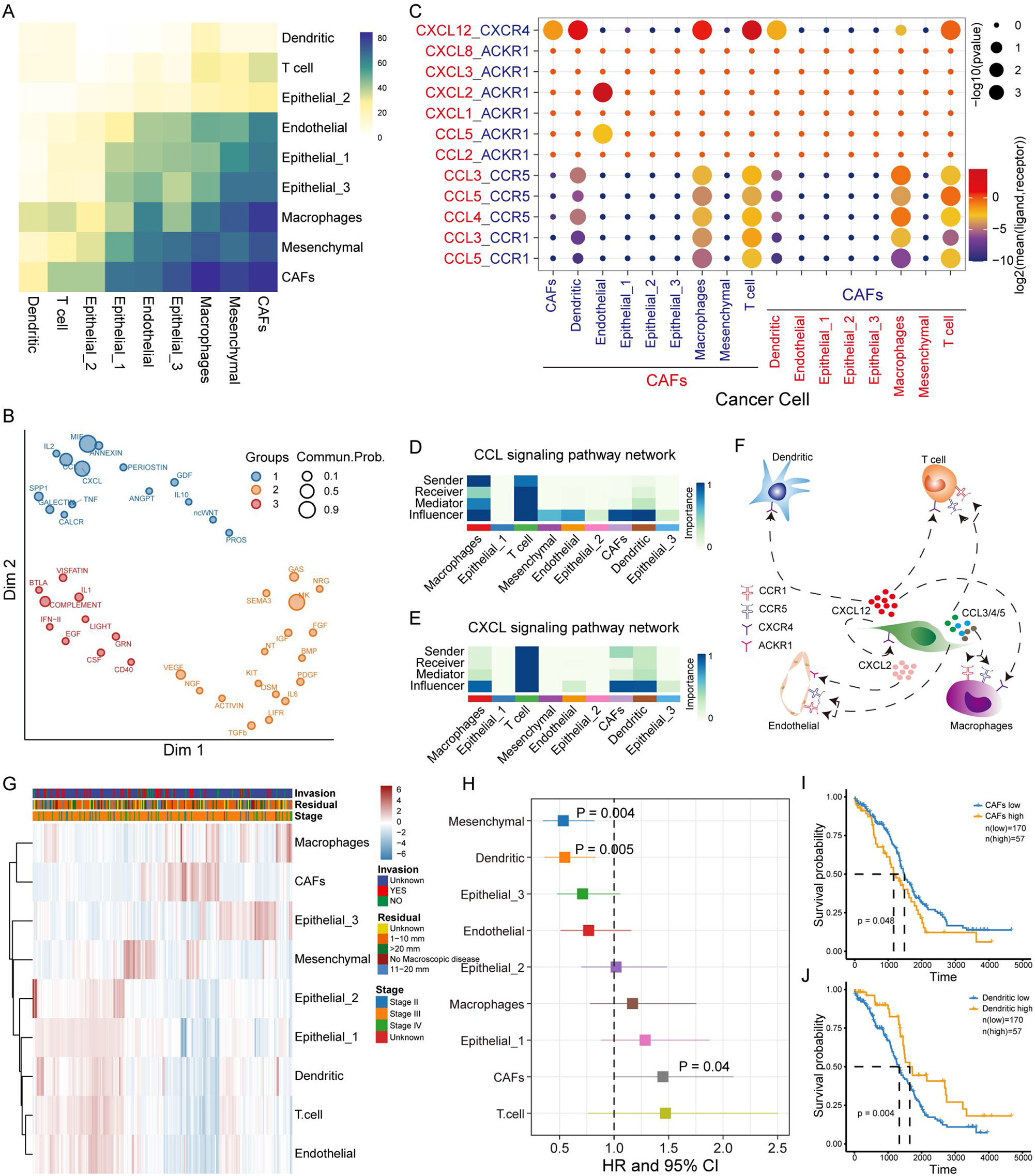
Cell-cell communication reveals the pro-metastatic effect of CAFs. (A) The heatmap shows the number of potential ligand-receptor pairs between cell populations predicted by CellChat. (B) Signaling pathways are clustered into 3 categories based on their function. (C) The bubble plot shows the ligand-receptor pairs of CXCL and CCL. (D) The amount of communication including sender, receiver, mediator and influencer between each cell type in the CXCL signaling pathway. (E) Same as in D but for CCL signaling pathway. (F) Predictive signaling factor regulatory network centered on CAFs. (G) Heatmap of cell abundance for each sample predicted based on bulk RNA-seq data by CIBERSORTx. Shown are row z-score. (H) COX risk regression to assess the association between relative cell abundance and patient survival from bulk RNA-seq. (I-J) Kaplan-Meier curves respond to patient survival in relation to CAFs and dendritic cells.

## Discussion

In this study, we re-analyzed a scRNA-seq datasets including 2794 high quality cells of EOC and characterized the physiological mechanisms reprograming in the TME. The malignant epithelial cells were defined by cell markers and CNVs and its individual difference and tumor heterogeneity was also observed. In TME of ovarian cancer, macrophages had global metabolic pathway activation and pathway score don’t have overall consistent between single-malignant cell and bulk tumor. Meanwhile, the activity of metabolic pathways contributes to the heterogeneity of malignant epithelial cells. Further, we simulated EMT trajectories using pseudo-time analysis, revealing that enhanced hypoxia, OXPHOS, and glycolysis together advance EMT process, which is also propelled by key factors identified by transcriptional regulation. The key signals of malignant epithelial cell metastasis based on cellular communication was identified, CAFs were found to be important signal transmitters involved in tumor metastasis and associated with OS of patients.

The relationship between energy metabolism and oxygen supply is intriguing. The positive correlation between hypoxia and glycolysis, which is consistent with the energy metabolism mechanism of Warburg effect (50). Hypoxia positively correlates with OXPHOS broke the phenomenon that OXPHOS is suppressed in tumors by comparing the expression of metabolic genes between bulk tumors and normal tissues (51, 52). The OXPHOS and glycolysis were important contributors to intratumor metabolic heterogeneity in malignant and stromal cells. Thus, the interaction between OXPHOS, glycolysis and hypoxia were highly dynamic in solid tumors. Then it is reasonable to infer that high OXPHOS activity consumes a large amount of oxygen causing cellular hypoxia, which activate hypoxia-induced factor (HIF). The quantitative relationship between OXPHOS and hypoxia is determined by the positive feedback formed with the HIF signaling pathway. Consistently, energy metabolic reprograming was found at single cell resolution were not identified in previous bulk tumor studies (53).

We used a reverse graph embedding method to simulate the developmental process of EMT and found the activity trajectory of energy metabolism and stemness. During the maturation of mesenchymal cells, the intensification of energy metabolism leads to increasing hypoxia. Several cell line experiments demonstrate that hypoxia induces EMT in tumor cells (54, 55), which is consistent with our findings. Our analysis revealed that key factors HGMA1 reduce the encoding of E-calmodulin creating an opportunity for epithelial cell metastasis, which could stimulate positive feedback regulation between increased energy metabolism and the HIF signaling pathway. To further evaluate the regulating roles of driver genes relating with EMT process, we considered the ceRNA competition mechanism and constructed a rational ceRNA regulatory network involving four lncRNA (two positive-correlation and two negative-correlation for EMT) and nine corresponding mRNA (seven positive-correlation and two negative-correlation for EMT, Supplementary Figure S13A). We found that GALNT1 expression is upregulated during EMT (Supplementary Figure S13B) and has been confirmed to promote tumor proliferation and metastasis in colorectal cancer (56). With the decreased expression of E-cadherin gradually, USP53 which up-regulated in expression (Supplementary Figure S13C) could promote apoptosis and inhibit glycolysis through FKBP51-AKT1 signaling (57). This finding is consistent with previous studies showing that cancer cells lacking E-calcineurin are more likely to undergo “apoptosis” although they are more easily subjected to aggressive growth (58). We found that the mechanism by which RHOBTB3 down-regulates HIFα protein levels is disrupted due to the high expression of RHOBTB3 accompanied by the accumulation of HIFα during EMT process (59) (Supplementary Figure S13D).

To summary, this study provides a global landscape of TME for EOC and characterized the physiological mechanism reprogramming at single-cell resolution. Although we analyzed the physiological mechanisms of EOC only in terms of metabolism and hallmark, this is also sufficient to explain tumor maintenance, proliferation and metastasis. All these conclusions in this study may provide more precise theoretical basis for the treatment strategy of EOC.

## Methods

### Data collection and pre-processing

The single-cell RNA-seq profiles for EOC was collected from the Gene Expression Omnibus (GEO (18); https://www.ncbi.nlm.nih.gov/gds) under accession numbers GSE118828. The profiles were taken from seven patients at different tissues and eighteen sequencing files were obtained, containing 3 064 single cells. After quality control (Supplementary Figure S1), 2 794 cells expression matrix was normalized using Scran (19) due to the low sequencing depth. The normalized matrix was used in principal component analysis (PCA) and clustering analysis using UMAP (20) implemented in the Seurat (21). Marker genes for specific cell types collected from CellMarker (22) (http://biocc.hrbmu.edu.cn/CellMarker/) and published literature (16) were used for the cell annotation. An independent single-cell RNA-seq profiles (GSE147082) including 9 885 cells from 6 EOC patients were used as validation dataset and were processed as above. Bulk RNA-seq (TCGA-OV) and microarray data (GSE26712) for EOC samples were collected for comparative analysis.

### The analysis of intercellular heterogeneity

The principal components analysis (PCA) was performed on the normalized RNA-seq profiles. For each gene in metabolic pathway and hallmark, we defined the weight of a gene as the sum of the absolute values of the loadings of this gene on the top PCs that explains at least 85% of the overall variation of the single-cell RNA-seq or bulk microarray profiles. Genes were sorted in descending order according to their weights and the Gene Set Enrichment Analysis (GSEA) (23) was used to identify metabolic pathways and hallmark enriched in the ranked genes with highest variability. Moreover, coefficient of variation (CV), standard deviation (SD), and information entropy were also used to interpret intercellular heterogeneity.

### Developmental trajectory analysis

To explore the process of epithelial mesenchymal transition (EMT), the epithelial cells, mesenchymal and CAFs were extracted for the construction of pseudo-time developmental trajectories using monocle (24) (v2.16.0). The break points and branches were marked in the developmental trajectory, where cells in the same segment of the trajectory were defined to have the same state.

### Construction of regulatory network for EMT-related genes

Pearson correlation analysis was used on the exploration of genes related to the pseudo-time. We defined genes with |P|>0.2 (25) as EMT-related genes. The expression variation direction of transcription factors (TFs) and corresponding target genes (TGs) included in EMT-related genes were supposed to be a regulatory pair in the pseudo-time process, that was *P*_*tf*_ x *P*_*tg*_ > 0. Human transcription factor and transcriptional regulatory interaction data were respectively retrieved from AnimalTFDB (26) 3.0 (http://bioinfo.life.hust.edu.cn/AnimalTFDB/), and TRRUST (27) v2 (www.grnpedia.org/trrust). Furthermore, we defined that the expression level was positively correlated for each lncRNA-mRNA pair due to the ceRNA competition mechanism. The lncRNA and mRNA relationship pairs in the ceRNA competition mechanism were acquired from the starBase (28) available at http://starbase.sysu.edu.cn/.

### Cell-cell communication analysis with CellChat

CellChat (29) is an R-based computational analysis tool developed by Suoqin Jin et al. that analyzes intercellular communication at the molecular level. All tumor cells that were defined as nine cell substes were used to explore the intercellular signaling interaction network using CellChat. Interaction pairs whose belong to signal transduction pathways and have *P*-values < 0.05 returned, were applied to explore the relationship between cell types in the tumor microenvironment.

### Connection to public datasets

Bulk RNA-seq and phenotype data of EOC organized by TCGA were downloaded from UCSC XENA (https://xenabrowser.net/datapages/). Then, the CIBERSORTx (30) algorithm, which is an analysis tool for abundance estimation of member cell types in mixed cell populations developed by Newman et al, was used to calculate the cellular fraction of each sample. Subsequently, bulk samples were categorized into high and low according to the three-fourths of the fraction of each cell type. The univariate Cox regression model (31) was used to estimate cell subsets in relation to overall survival (OS) of EOC patients. Kaplan-Meier analysis (32) was performed to estimate the OS curves of samples.

### Reagents and cell cultures

Antibodies against EGR1 and EMT-relative kit (E-cadherin, N-cadherin, and slug) were purchased from Cell Signaling Technology (Danvers, MA). Two human ovarian cancer cell lines (SKOV3 and A2780) were used in our study. The SKOV-3 and A2780 cell lines were purchased from American Type Culture Collection (ATCC, South San Francisco, CA). Cells were cultured in Dulbecco modified Eagle medium (DMEM; Life Technologies, Grand Island, NY) with 10% fetal bovine serum (Life Technologies) and were maintained in a humidified 5% CO2 incubator at 37°C.

### Wound healing assay

Cells transfected with si-NC, si-EGR1 were seeded into six-well culture plates with serum-containing medium and were cultured until the cell density reached 90%-95% confluence. An artificial homogeneous wound was created by scratching the monolayer with a sterile 200 μL pipette tip. After scratching, the cells were washed with PBS, and then the cells were cultured with serum-free DMEM media for 48 hours. Images of cells migrating into the wound were captured at 0 and 48 hours using a microscope (EVOS, USA). The assay was performed in triplicate.

### Cell Migration Assay

Cell migration and invasion were determined by the trans-well assay. Polycarbonate trans-well filters with 8.0-μm pores were inserted over the lower chambers. A total of 5 × 104 cells suspended in 2% FBS were plated on the insert chamber supplemented with 600 μl of 10% FBS medium. After 12 h, cells were fixed and permeabilized in propyl alcohol for 15 min and then stained with 5% crystal violet stain overnight at 4°C. The number and morphology of cells were observed under an inverted microscope.

### Western blotting

Extraction of the total protein was finished with RIPA kit. The protein was separated with SDS-PAGE (12.5%), moved to NC membrane, and later blocked with 5% non-fat milk for two hours at room temperature (Thermo fisher). Then, the membrane was incubated with primary antibodies to E-cadherin, slug, N-cadherin, HSPG2, FAS and EGR1 at 4°C overnight. After that, the membrane was incubated with a secondary antibody to HRP-labeled Beyotime, Beijing. Blots were viewed with electrochemiluminescence (ECL) chromogenic substrate.

### Statistical analysis

All statistical analyses and graph generation were performed in R (version 4.0.2).

## Ethics approval and consent to participate

Not applicable.

## Consent for publication

Not applicable.

## Availability of data and materials

All the data used in the analysis can be obtained from TCGA, ENCODE, and GEO.

## Authors’ contributions

YZ, SS and YH designed the study, performed analysis, wrote and revised the manuscript. YQ and YD revised the manuscript. MX, RX, CK and HC helped with data collection. XW and QL proofread the manuscript and linguistic touch-ups. GZ, PW and QG share the senior authorship of this study. The authors read and approved the final manuscript.

## Acknowledgements

The authors gratefully thank the TCGA, ENCODE, and GEO for providing data for this work.

